## Supplementary material for "Two neuropeptides that promote blood feeding in *Anopheles stephensi* mosquitoes": Supp sheet1_GLMM analysis

### GLMM analysis of feeding behaviours in *An. stephensi* females (figure 1D, E)

*Data from D0 (day of emergence) were excluded from the GLMM analysis because newly emerged females show no feeding behaviour on this day, resulting in no variance in the response variable to model.*

#### Model 1: Whether the experimental groups differ in blood-feeding behaviour across age?

Random variables: BioID (replicate identity), larval.batch (larval batch)

Fixed variables: Day(age), Group (co-housed vs virgin)

```
model1 <- glmer(Bloodfed_binary ~ Day * Group + (1|BioID) + (1|larval.batch),
        data = mosquito_data_filtered,
        family = binomial)
```

**Model summary:**

| Characteristic | Odds | Probability (%) | OR[1] | 95% CI | p-value |
|---|---|---|---|---|---|
| **Day** | | | | | |
| D1 | 1.41 | 58.5 | — | — | |
| D2 | 2.67 | 72.8 | 1.90** | 1.29, 2.81 | **0.001** |
| D3 | 4.87 | 83.0 | 3.47*** | 2.25, 5.35 | **<0.001** |
| D4 | 6.35 | 86.4 | 4.52*** | 2.86, 7.13 | **<0.001** |
| D5 | 4.64 | 82.3 | 3.30*** | 2.15, 5.08 | **<0.001** |
| **Group** | | | | | |
| co-housed | 1.41 | 58.5 | — | — | |
| virgin | 1.20 | 54.5 | 0.86 | 0.59, 1.23 | 0.403 |
| **Day * Group** | | | | | |
| D2 * virgin | 2.44 | 70.9 | 1.74 | 0.99, 3.05 | 0.053 |
| D3 * virgin | 0.95 | 48.7 | 0.68 | 0.38, 1.21 | 0.189 |
| D4 * virgin | 0.56 | 35.9 | 0.40** | 0.22, 0.72 | **0.002** |
| D5 * virgin | 0.90 | 47.4 | 0.64 | 0.36, 1.14 | 0.130 |

[1] *p<0.05; **p<0.01; ***p<0.001

Abbreviations: CI = Confidence Interval, OR = Odds Ratio

### Post-hoc analysis via estimated marginal means (EM means)

#### 1) Age-effects within each group (p-value adjustment: Tukey's method)

Group = co-housed:

| contrast | estimate | SE | df | z.ratio | p.value |
|---|---|---|---|---|---|
| D1 - D2 | -0.6423 | 0.199 | Inf | -3.234 | 0.0107 |
| D1 - D3 | -1.2432 | 0.221 | Inf | -5.614 | <.0001 |
| D1 - D4 | -1.5085 | 0.233 | Inf | -6.479 | <.0001 |
| D1 - D5 | -1.1946 | 0.220 | Inf | -5.429 | <.0001 |
| D2 - D3 | -0.6009 | 0.228 | Inf | -2.638 | 0.0637 |
| D2 - D4 | -0.8662 | 0.242 | Inf | -3.577 | 0.0032 |
| D2 - D5 | -0.5523 | 0.229 | Inf | -2.416 | 0.1111 |
| D3 - D4 | -0.2653 | 0.261 | Inf | -1.018 | 0.8473 |
| D3 - D5 | 0.0485 | 0.248 | Inf | 0.196 | 0.9997 |
| D4 - D5 | 0.3138 | 0.262 | Inf | 1.199 | 0.7521 |

Group = virgin:

| contrast | estimate | SE | df | z.ratio | p.value |
|---|---|---|---|---|---|
| D1 - D2 | -1.1958 | 0.210 | Inf | -5.697 | <.0001 |
| D1 - D3 | -0.8564 | 0.201 | Inf | -4.262 | 0.0002 |
| D1 - D4 | -0.5857 | 0.192 | Inf | -3.050 | 0.0194 |
| D1 - D5 | -0.7529 | 0.199 | Inf | -3.791 | 0.0014 |
| D2 - D3 | 0.3394 | 0.220 | Inf | 1.542 | 0.5353 |
| D2 - D4 | 0.6100 | 0.216 | Inf | 2.826 | 0.0379 |
| D2 - D5 | 0.4429 | 0.221 | Inf | 2.007 | 0.2625 |
| D3 - D4 | 0.2706 | 0.207 | Inf | 1.308 | 0.6859 |
| D3 - D5 | 0.1035 | 0.211 | Inf | 0.491 | 0.9882 |
| D4 - D5 | -0.1671 | 0.207 | Inf | -0.806 | 0.9288 |

Results are given on the log odds ratio (not the response) scale.
P value adjustment: tukey method for comparing a family of 5 estimates

### 2) Pair-wise comparison at each day (p-value adjustment: Bonferroni method)

```
Day = D1:
 contrast               estimate    SE  df z.ratio p.value
 (co-housed) - virgin      0.156 0.186 Inf   0.836  0.4034

Day = D2:
 contrast               estimate    SE  df z.ratio p.value
 (co-housed) - virgin     -0.398 0.217 Inf  -1.830  0.0672

Day = D3:
 contrast               estimate    SE  df z.ratio p.value
 (co-housed) - virgin      0.542 0.228 Inf   2.376  0.0175

Day = D4:
 contrast               estimate    SE  df z.ratio p.value
 (co-housed) - virgin      1.078 0.236 Inf   4.570  <.0001

Day = D5:
 contrast               estimate    SE  df z.ratio p.value
 (co-housed) - virgin      0.597 0.225 Inf   2.656  0.0079

Results are given on the log odds ratio (not the response) scale.
```

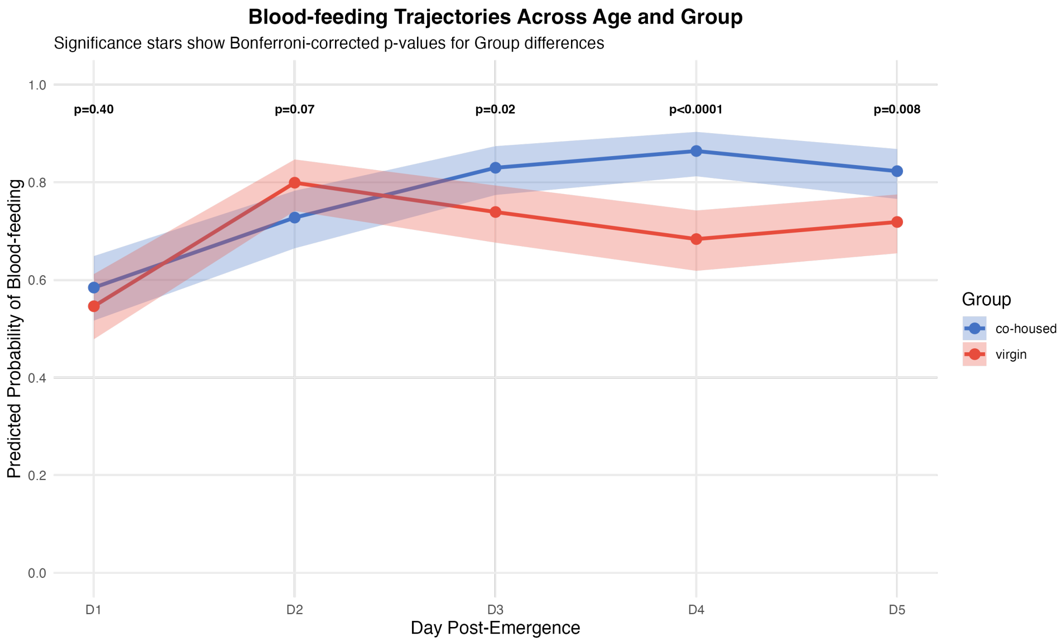


**<u>Conclusions:</u>** Blood-feeding probability of co-housed females increases with age. This trend is weaker in virgins.

Mated females feed significantly higher than virgins as they age (D3-D5), the differences being most pronounced on D4.

**Model 2: Effect of mating on the blood-feeding behaviour of co-housed females**

Random variables: BioID (replicate identity)

Fixed variables: Day(age), Mating status (yes or no)

```
model2 <- glmer(Bloodfed_binary ~ Day * Mating + (1|BioID),
            data = mated_only,
            family = binomial)
```

**Model summary:**

| Characteristic | Odds | Probability (%) | OR[1] | 95% CI | p-value |
|---|---|---|---|---|---|
| **Day** | | | | | |
| D1 | 1.37 | 57.8 | — | — | |
| D2 | 2.14 | 68.2 | 1.56 | 1.00, 2.46 | 0.052 |
| D3 | 2.23 | 69.0 | 1.63 | 0.87, 3.06 | 0.130 |
| D4 | 4.23 | 80.9 | 3.09* | 1.21, 7.88 | **0.018** |
| D5 | 1.54 | 60.6 | 1.12 | 0.18, 6.89 | 0.902 |
| **Mating** | | | | | |
| n | | | — | — | |
| y | 2.63 | 72.5 | 1.92 | 0.49, 7.46 | 0.347 |
| **Day * Mating** | | | | | |
| D2 * y | 1.22 | 55.0 | 0.89 | 0.20, 3.92 | 0.878 |
| D3 * y | 2.18 | 68.6 | 1.59 | 0.34, 7.42 | 0.555 |
| D4 * y | 1.14 | 53.3 | 0.83 | 0.16, 4.42 | 0.827 |
| D5 * y | 2.28 | 69.5 | 1.67 | 0.17, 16.2 | 0.660 |

[1] *p<0.05; **p<0.01; ***p<0.001

Abbreviations: CI = Confidence Interval, OR = Odds Ratio

### Post-hoc analysis via estimated marginal means (EM means)

#### Effect of mating (y) on each day (p-value adjustment: Bonferroni method)

```
Day = D1:
 contrast estimate    SE  df z.ratio p.value
 y - n       0.651 0.693 Inf   0.939  0.3475

Day = D2:
 contrast estimate    SE  df z.ratio p.value
 y - n       0.535 0.301 Inf   1.777  0.0756

Day = D3:
 contrast estimate    SE  df z.ratio p.value
 y - n       1.115 0.368 Inf   3.027  0.0025

Day = D4:
 contrast estimate    SE  df z.ratio p.value
 y - n       0.464 0.504 Inf   0.920  0.3575

Day = D5:
 contrast estimate    SE  df z.ratio p.value
 y - n       1.162 0.934 Inf   1.245  0.2133

Results are given on the log odds ratio (not the response) scale.
```

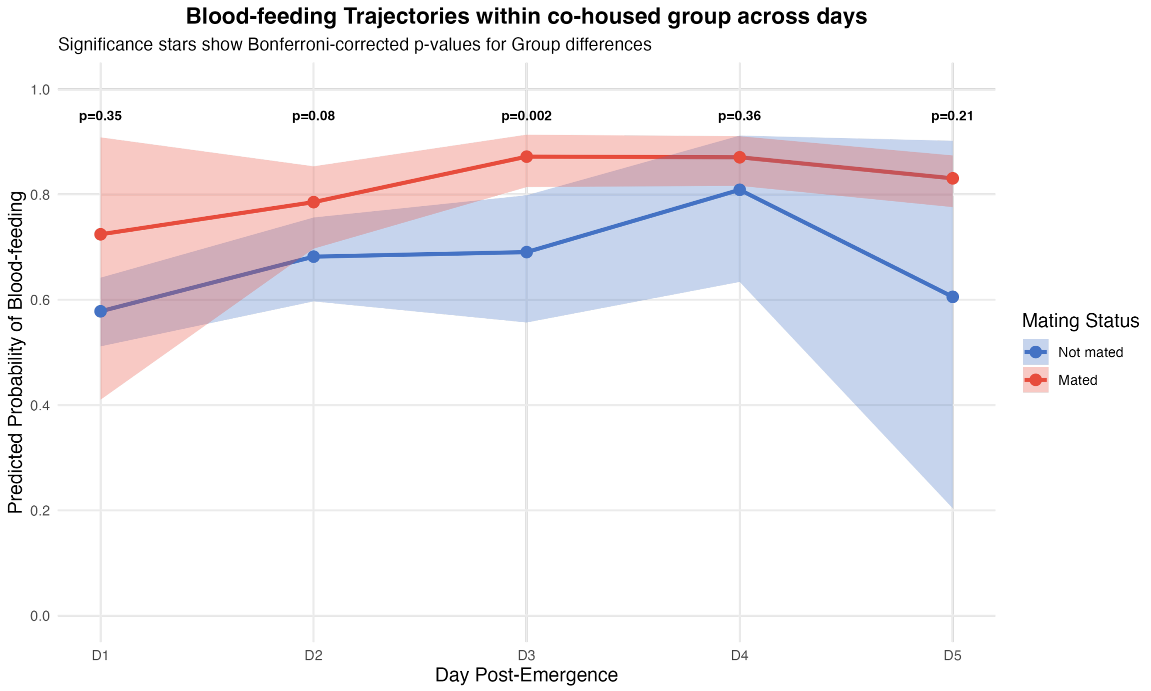


**Conclusions:** Within co-housed females, mating status shows only a brief, significant effect on blood-feeding on D3. The pronounced differences observed in the blood-feeding behaviour of virgin and co-housed females on Days 4-5 (model 1) appear to be mediated by male presence rather than mating per se, as mated and unmated females within the co-housed group show no significant differences during these time points.

**Model 3: Whether the experimental groups differ in their second-blood meal, across age?**

Random variables: BioID (replicate identity)

Fixed variables: Day(age), Group (co-housed vs virgin)

```
model3 <- glmer(Bloodfed_binary ~ Day * Group + (1|BioID),
            data = mosquito_data,
            family = binomial)
```

**Model summary:**

| Characteristic | Odds | Probability (%) | OR[1] | 95% CI | p-value |
|---|---|---|---|---|---|
| **Day** | | | | | |
| DB1 | 0.51 | 33.8 | — | — | |
| DB2 | 0.21 | 17.4 | 0.41*** | 0.29, 0.60 | **<0.001** |
| DB3 | 0.65 | 39.4 | 1.28 | 0.92, 1.78 | 0.143 |
| DB4 | 0.40 | 28.6 | 0.78 | 0.55, 1.10 | 0.154 |
| **Group** | | | | | |
| co-housed | 0.51 | 33.8 | — | — | |
| virgin | 4.35 | 81.3 | 8.58*** | 5.81, 12.7 | **<0.001** |
| **Day * Group** | | | | | |
| DB2 * virgin | 1.47 | 59.5 | 2.90*** | 1.67, 5.02 | **<0.001** |
| DB3 * virgin | 0.47 | 32.0 | 0.92 | 0.54, 1.55 | 0.748 |
| DB4 * virgin | 0.97 | 49.2 | 1.91* | 1.11, 3.28 | **0.020** |

[1] *p<0.05; **p<0.01; ***p<0.001

Abbreviations: CI = Confidence Interval, OR = Odds Ratio

(note: PBM abbreviated as “B”)

**Post-hoc analysis via estimated marginal means (EM means)**

1) **Age-effects within each group (p-value adjustment: Tukey's method)**

```
Group = co-housed:
 contrast  estimate    SE  df z.ratio p.value
 DB1 - DB2   0.8849 0.188 Inf   4.703  <.0001
 DB1 - DB3  -0.2477 0.169 Inf  -1.467  0.4579
 DB1 - DB4   0.2499 0.175 Inf   1.424  0.4841
 DB2 - DB3  -1.1326 0.174 Inf  -6.498  <.0001
 DB2 - DB4  -0.6350 0.180 Inf  -3.524  0.0024
 DB3 - DB4   0.4976 0.161 Inf   3.092  0.0107

Group = virgin:
 contrast  estimate    SE  df z.ratio p.value
 DB1 - DB2  -0.1790 0.210 Inf  -0.852  0.8296
 DB1 - DB3  -0.1619 0.210 Inf  -0.773  0.8667
 DB1 - DB4  -0.3952 0.217 Inf  -1.822  0.2629
 DB2 - DB3   0.0171 0.202 Inf   0.085  0.9998
 DB2 - DB4  -0.2162 0.210 Inf  -1.032  0.7308
 DB3 - DB4  -0.2332 0.209 Inf  -1.117  0.6792

Results are given on the log odds ratio (not the response) scale.
P value adjustment: tukey method for comparing a family of 4 estimates
```

2) **Pair-wise comparison at each day (p-value adjustment: Bonferroni method)**

```
Day = DB1:
 contrast             estimate    SE  df z.ratio p.value
 (co-housed) - virgin    -2.15 0.199 Inf -10.800  <.0001

Day = DB2:
 contrast             estimate    SE  df z.ratio p.value
 (co-housed) - virgin    -3.21 0.199 Inf -16.129  <.0001

Day = DB3:
 contrast             estimate    SE  df z.ratio p.value
 (co-housed) - virgin    -2.06 0.180 Inf -11.492  <.0001

Day = DB4:
 contrast             estimate    SE  df z.ratio p.value
 (co-housed) - virgin    -2.79 0.194 Inf -14.397  <.0001

Results are given on the log odds ratio (not the response) scale.
```

(note: PBM abbreviated as "B")

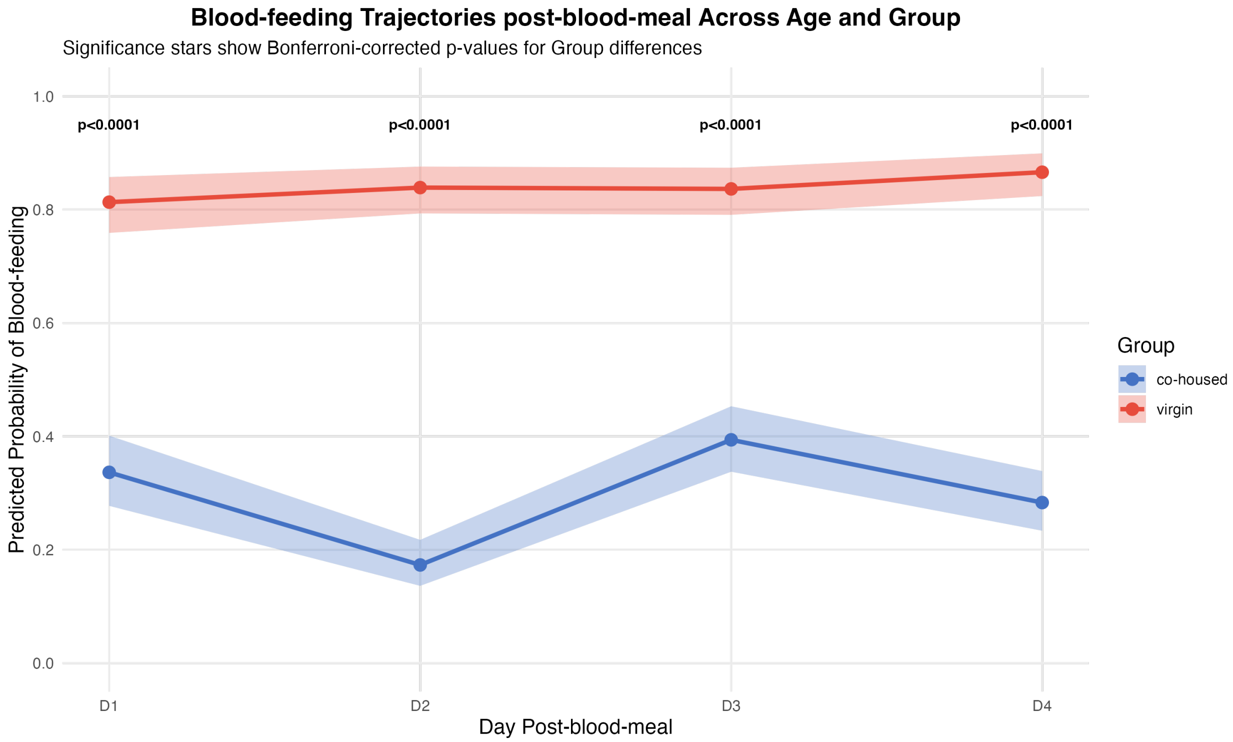


**<u>Conclusions:</u>** Co-housed and virgin females show markedly different appetites, post-blood meal. Co-housed females (assumed mated), show a strong sustained suppression, while virgins continue to feed with high propensity, at least up to 4 days after a replete blood meal.

**Model 4: Whether the experimental groups differ in sugar-feeding behaviour across age?**

Random variables: BioID (replicate identity)

Fixed variables: Day(age), Group (co-housed vs virgin)

```
model4 <- glmer(Sugar.fed_binary ~ Day * Group + (1|BioID),
            data = mosquito_data_filtered,
            family = binomial)
```

**Model summary:**

| Characteristic | Odds | Probability (%) | OR[1] | 95% CI | p-value |
|---|---|---|---|---|---|
| **Day** | | | | | |
| D1 | 0.41 | 29.1 | — | — | |
| D2 | 12.37 | 92.5 | 30.1*** | 15.6, 57.9 | **<0.001** |
| D3 | 17.88 | 94.7 | 43.4*** | 20.6, 91.8 | **<0.001** |
| D4 | 33.53 | 97.1 | 81.5*** | 31.6, 210 | **<0.001** |
| D5 | 28.03 | 96.6 | 68.1*** | 28.3, 164 | **<0.001** |
| **Group** | | | | | |
| co-housed | 0.41 | 29.1 | — | — | |
| virgin | 0.88 | 46.8 | 2.13*** | 1.38, 3.27 | **<0.001** |
| **Day * Group** | | | | | |
| D2 * virgin | 0.21 | 17.4 | 0.51 | 0.21, 1.23 | 0.134 |
| D3 * virgin | 0.09 | 8.3 | 0.21*** | 0.09, 0.52 | **<0.001** |
| D4 * virgin | 0.16 | 13.8 | 0.39 | 0.12, 1.30 | 0.126 |
| D5 * virgin | 0.15 | 13.0 | 0.37 | 0.12, 1.13 | 0.080 |

[1] *p<0.05; **p<0.01; ***p<0.001

Abbreviations: CI = Confidence Interval, OR = Odds Ratio

### Post-hoc analysis via estimated marginal means (EM means)

#### 1) Age-effects within each group (p-value adjustment: Tukey's method)

Group = co-housed:

| contrast | estimate | SE | df | z.ratio | p.value |
|---|---|---|---|---|---|
| D1 - D2 | -3.403 | 0.334 | Inf | -10.185 | <.0001 |
| D1 - D3 | -3.771 | 0.382 | Inf | -9.884 | <.0001 |
| D1 - D4 | -4.400 | 0.484 | Inf | -9.093 | <.0001 |
| D1 - D5 | -4.221 | 0.448 | Inf | -9.421 | <.0001 |
| D2 - D3 | -0.368 | 0.448 | Inf | -0.823 | 0.9238 |
| D2 - D4 | -0.997 | 0.537 | Inf | -1.855 | 0.3419 |
| D2 - D5 | -0.818 | 0.506 | Inf | -1.617 | 0.4861 |
| D3 - D4 | -0.629 | 0.568 | Inf | -1.106 | 0.8032 |
| D3 - D5 | -0.449 | 0.538 | Inf | -0.835 | 0.9198 |
| D4 - D5 | 0.179 | 0.615 | Inf | 0.292 | 0.9984 |

Group = virgin:

| contrast | estimate | SE | df | z.ratio | p.value |
|---|---|---|---|---|---|
| D1 - D2 | -2.737 | 0.296 | Inf | -9.238 | <.0001 |
| D1 - D3 | -2.224 | 0.256 | Inf | -8.685 | <.0001 |
| D1 - D4 | -3.454 | 0.387 | Inf | -8.916 | <.0001 |
| D1 - D5 | -3.225 | 0.354 | Inf | -9.114 | <.0001 |
| D2 - D3 | 0.514 | 0.337 | Inf | 1.525 | 0.5461 |
| D2 - D4 | -0.717 | 0.445 | Inf | -1.613 | 0.4887 |
| D2 - D5 | -0.488 | 0.416 | Inf | -1.173 | 0.7668 |
| D3 - D4 | -1.231 | 0.419 | Inf | -2.937 | 0.0274 |
| D3 - D5 | -1.001 | 0.388 | Inf | -2.579 | 0.0743 |
| D4 - D5 | 0.230 | 0.485 | Inf | 0.474 | 0.9897 |

Results are given on the log odds ratio (not the response) scale.
P value adjustment: tukey method for comparing a family of 5 estimates

#### 2) Pair-wise comparison at each day (p-value adjustment: Bonferroni method)

Day = D1:

| contrast | estimate | SE | df | z.ratio | p.value |
|---|---|---|---|---|---|
| (co-housed) - virgin | -0.7546 | 0.219 | Inf | -3.441 | 0.0006 |

Day = D2:

| contrast | estimate | SE | df | z.ratio | p.value |
|---|---|---|---|---|---|
| (co-housed) - virgin | -0.0887 | 0.392 | Inf | -0.226 | 0.8210 |

Day = D3:

| contrast | estimate | SE | df | z.ratio | p.value |
|---|---|---|---|---|---|
| (co-housed) - virgin | 0.7932 | 0.407 | Inf | 1.950 | 0.0512 |

Day = D4:

| contrast | estimate | SE | df | z.ratio | p.value |
|---|---|---|---|---|---|
| (co-housed) - virgin | 0.1911 | 0.582 | Inf | 0.328 | 0.7426 |

Day = D5:

| contrast | estimate | SE | df | z.ratio | p.value |
|---|---|---|---|---|---|
| (co-housed) - virgin | 0.2413 | 0.529 | Inf | 0.456 | 0.6485 |

Results are given on the log odds ratio (not the response) scale.

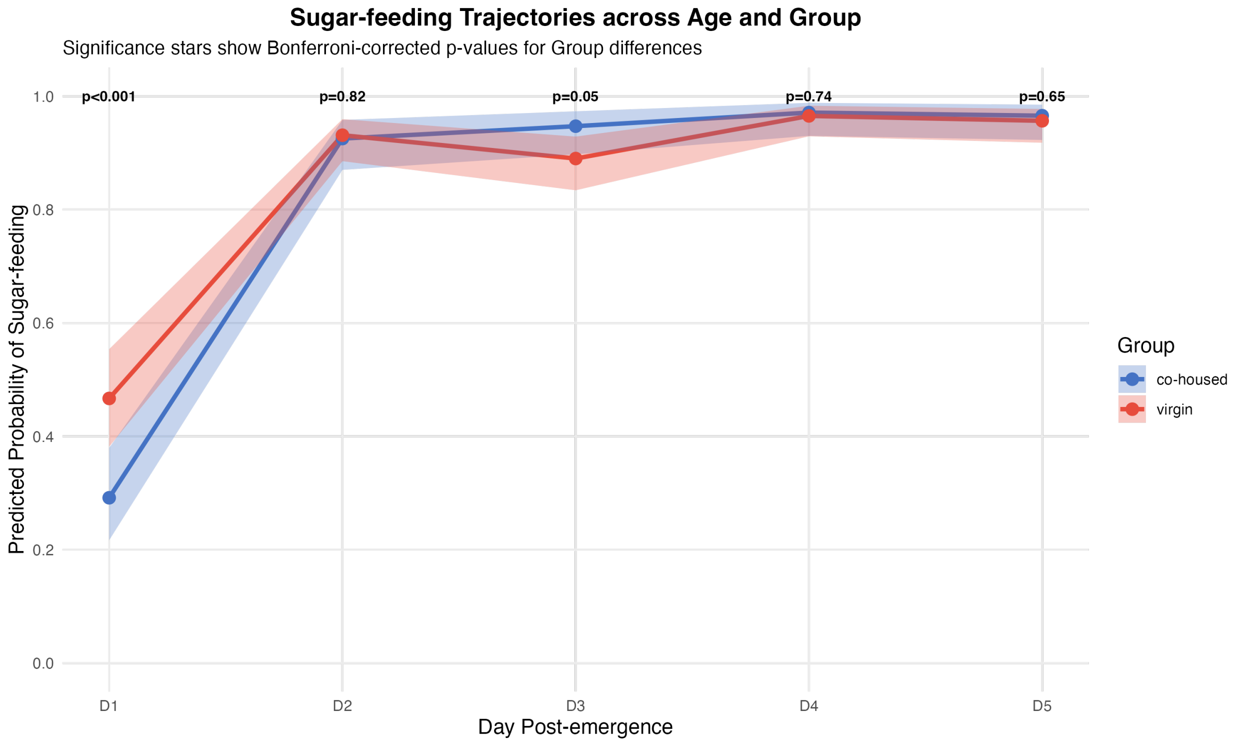


**<u>Conclusions:</u>** Both virgins and co-housed females show a sharp increase in their sugar-appetite from D1 to D2, followed by stabilization through D5.

Sugar-feeding behaviours are largely comparable between groups, across days. Virgin females demonstrated significantly higher initial feeding probability compared to co-housed females (D1). This difference was eliminated by D2 and remained non-significant thereafter.

### Model 5: Effect of mating on the sugar-feeding behaviour of co-housed females

Random variables: BioID (replicate identity)

Fixed variables: Day(age), Mating status (yes or no)

```
model5 <- glmer(Sugar.fed_binary ~ Day * Mating + (1|BioID),
        data = mated_only,
        family = binomial)
```

**Model summary:**

| Characteristic | Odds | Probability (%) | OR[1] | 95% CI | p-value |
|---|---|---|---|---|---|
| **Day** | | | | | |
| D1 | 0.42 | 29.6 | — | — | |
| D2 | 7.75 | 88.6 | 18.4*** | 8.19, 41.5 | **<0.001** |
| D3 | 2.98 | 74.9 | 7.08*** | 2.64, 19.0 | **<0.001** |
| D4 | 6.48 | 86.6 | 15.4*** | 4.32, 55.1 | **<0.001** |
| D5 | 13511668.03 | 100.0 | 32,155,341 | 0.00, 458,391,919,179,031,447,577,326,110,191,003,156,070,149,023,901,469,250,371,113,818,366,346,793,494,511,616 | 0.841 |
| **Mating** | | | | | |
| n | | | — | — | |
| y | 1.70 | 63.0 | 4.05 | 0.92, 17.9 | 0.065 |
| **Day * Mating** | | | | | |
| D2 * y | 0.28 | 21.9 | 0.67 | 0.10, 4.44 | 0.682 |
| D3 * y | 1.76 | 63.8 | 4.18 | 0.52, 33.8 | 0.180 |
| D4 * y | 1.24 | 55.4 | 2.95 | 0.27, 31.7 | 0.373 |
| D5 * y | 0.00 | 0.0 | 0.00 | 0.00, 7,248,915,324,456,836,893,750,989,915,264,405,095,298,755,374,445,726,709,201,567,219,712 | 0.866 |

[1] *p<0.05; **p<0.01; ***p<0.001

Abbreviations: CI = Confidence Interval, OR = Odds Ratio

### Post-hoc analysis via estimated marginal means (EM means)

#### Effect of mating (yes) on each day (p-value adjustment: Bonferroni method)

```
Day = D1:
 contrast estimate    SE  df z.ratio p.value
 y - n        1.40 0.757 Inf   1.846  0.0648

Day = D2:
 contrast estimate    SE  df z.ratio p.value
 y - n        1.00 0.596 Inf   1.684  0.0921

Day = D3:
 contrast estimate    SE  df z.ratio p.value
 y - n        2.83 0.754 Inf   3.753  0.0002

Day = D4:
 contrast estimate    SE  df z.ratio p.value
 y - n        2.48 0.946 Inf   2.620  0.0088

Day = D5:
 contrast estimate    SE  df z.ratio p.value
 y - n      -13.08 85.900 Inf  -0.152  0.8790

Results are given on the log odds ratio (not the response) scale.
```

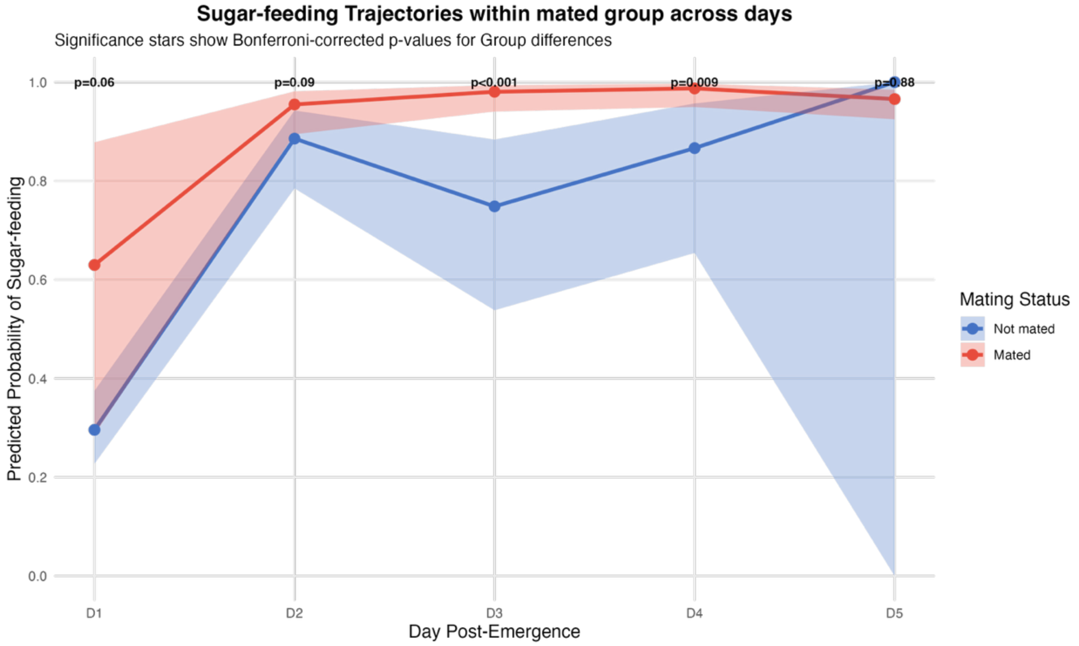


**Conclusions:** Mating status shows only a brief, significant effect on sugar-feeding on D3 and D4 within co-housed females: mating positively influences sugar-feeding on these days. However, it is interesting to note that no differences were observed in the feeding behaviour of virgins and co-housed females (model 4) on these days. As mating sharply increases from D2 to D4, it possibly creates maximum differences on these days within the co-housed group, which are then cancelled out when averaged together.

**Model 6: Whether the experimental groups differ in their sugar-feeding behaviour, post-blood meal, across age?**

Random variables: BioID (replicate identity)

Fixed variables: Day(age), Group (co-housed vs virgin)

```
model6 <- glmer(Sugar.fed_binary ~ Day * Group + (1|BioID),
        data = mosquito_data,
        family = binomial)
```

**Model summary:**

| Characteristic | Odds | Probability (%) | OR[1] | 95% CI | p-value |
|---|---|---|---|---|---|
| **Day** | | | | | |
| DB1 | 0.02 | 2.0 | — | — | |
| DB2 | 0.05 | 4.8 | 3.13 | 0.83, 11.8 | 0.093 |
| DB3 | 10.11 | 91.0 | 631*** | 180, 2,219 | **<0.001** |
| DB4 | 389103395.39 | 100.0 | 24,298,538,607 | 0.00, 347,082,619,877,364,307,044,218,276,151,296 | 0.358 |
| **Group** | | | | | |
| co-housed | 0.02 | 2.0 | — | — | |
| virgin | 0.22 | 18.0 | 13.6*** | 4.12, 45.2 | **<0.001** |
| **Day * Group** | | | | | |
| DB2 * virgin | 0.12 | 10.7 | 7.21** | 1.74, 29.8 | **0.006** |
| DB3 * virgin | 0.01 | 1.0 | 0.43 | 0.08, 2.17 | 0.306 |
| DB4 * virgin | 0.00 | 0.0 | 0.00 | 0.00, 124,369,836,581,060 | 0.476 |

[1] *p<0.05; **p<0.01; ***p<0.001

Abbreviations: CI = Confidence Interval, OR = Odds Ratio

(note: PBM abbreviated as “B”)

**Post-hoc analysis via estimated marginal means (EM means)**

1) **Age-effects within each group (p-value adjustment: Tukey's method)**

```
Group = co-housed:
 contrast  estimate     SE  df z.ratio p.value
 DB1 - DB2    -1.14  0.678 Inf  -1.680  0.3343
 DB1 - DB3    -6.45  0.641 Inf -10.055  <.0001
 DB1 - DB4   -23.91 26.000 Inf  -0.919  0.7948
 DB2 - DB3    -5.31  0.433 Inf -12.246  <.0001
 DB2 - DB4   -22.77 26.000 Inf  -0.875  0.8179
 DB3 - DB4   -17.47 26.000 Inf  -0.671  0.9081

Group = virgin:
 contrast  estimate     SE  df z.ratio p.value
 DB1 - DB2    -3.12  0.257 Inf -12.141  <.0001
 DB1 - DB3    -5.60  0.539 Inf -10.387  <.0001
 DB1 - DB4    -5.35  0.491 Inf -10.896  <.0001
 DB2 - DB3    -2.49  0.538 Inf  -4.623  <.0001
 DB2 - DB4    -2.24  0.489 Inf  -4.569  <.0001
 DB3 - DB4     0.25  0.679 Inf   0.368  0.9830

Results are given on the log odds ratio (not the response) scale.
P value adjustment: tukey method for comparing a family of 4 estimates
```

2) **Pair-wise comparison at each day (p-value adjustment: Bonferroni method)**

```
Day = DB1:
 contrast               estimate     SE  df z.ratio p.value
 (co-housed) - virgin      -2.61  0.611 Inf  -4.275  <.0001

Day = DB2:
 contrast               estimate     SE  df z.ratio p.value
 (co-housed) - virgin      -4.59  0.398 Inf -11.523  <.0001

Day = DB3:
 contrast               estimate     SE  df z.ratio p.value
 (co-housed) - virgin      -1.77  0.562 Inf  -3.141  0.0017

Day = DB4:
 contrast               estimate     SE  df z.ratio p.value
 (co-housed) - virgin      15.95 26.000 Inf   0.613  0.5401

Results are given on the log odds ratio (not the response) scale.
```

(note: PBM abbreviated as "B")

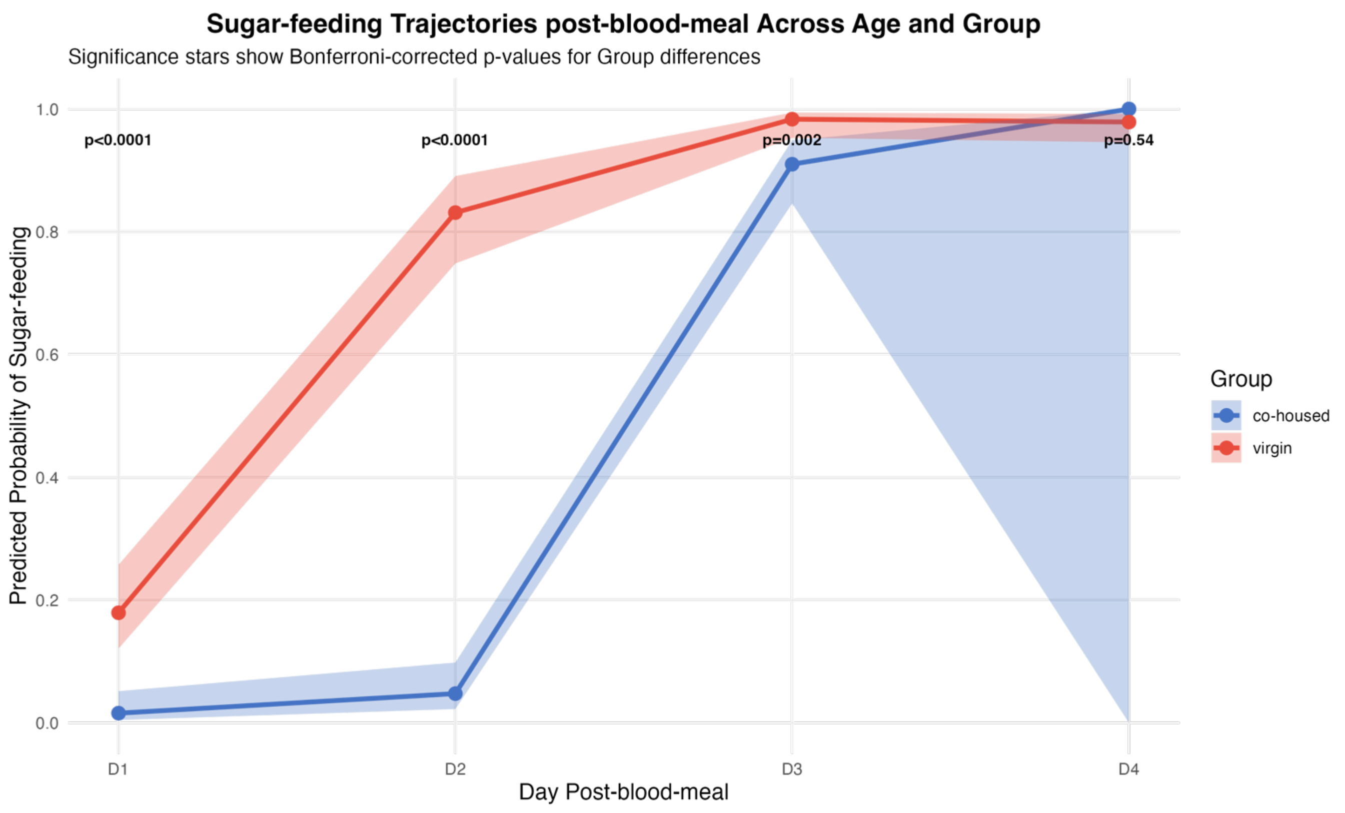


Conclusions: Both virgins and co-housed females (assumed mated) show a repressed appetite for sugar immediately after a blood meal (D1 post-blood meal). This repression is significantly stronger in mated females. Virgins start to regain their appetite by D2 post-blood meal, while the mated females continue to suppress it. By D3 post-blood meal, the appetite is restored for both groups.
